# A comparison of Alberta’s Infant Motor Scale and Brazelton Neonatal Behavior Assessment Scale

**DOI:** 10.1101/462572

**Authors:** Priyantha Perera, Pavithra Godamunne, Sumal Nandasena, Meghan Tipre, Udaya Wimalasiri, Mark Reader, Thulani Beddage, Tamika Smith, Anuradhani Kasturiratne, Nalini Sathiakumar, Rajitha Wickremasinghe

## Abstract

**Background and aims:** The Neonatal Behavior Assessment Scale (NBAS) was introduced by Brazelton is a useful tool to assess neurodevelopment in newborn in research setting. However, it is complex, time-consuming and requires specialized training for administration and is therefore difficult tool to use in routine clinical settings. The Alberta Infant Motor Scale (AIMS) is a simple tool to assess of motor maturation of a child up to 18 months and can be administered with minimal training for the health care professionals. The study goal of this study was to evaluate the reliability and validity of the AIMS in comparison with the NBAS in assessing motor maturity at birth.

**Methods:** We administered the AIMS and NBAS to a total of 66 newborn babies delivered in three obstetric units at the Colombo North Teaching Hospital in Ragama, Sri Lanka. The subjects were selected from a sample of 545 newborn babies in an ongoing birth-cohort study evaluating effects of prenatal exposure to biomass smoke and infant neurodevelopment. Trained research assistants administered first the NBAS, followed by the AIMS one-hour later. Univariate and bivariate statistics were used to compare the two scales.

**Results:** Irrespective of maturity, sex, birth weight or socio-demographic characteristics, all babies had scored on the 75th percentile in the AIMS. In the NBAS, there was a significant variation in the Brazelton motor score scaled to 100. Low birth weight babies showed a narrower variation in the NABS score. None of the scales indicated a motor deficit in any of the children.

**Conclusion:** NBAS identifies subtle differences in motor maturity of full term babies that the AIMS fails to detect

## Introduction and background

Assessment of neurodevelopment is an essential component of paediatric care. However, assessment of neurodevelopment of a newborn is performed rarely in clinical practice. In 1955, a Neonatal Behavior Assessment Scale (NBAS) was introduced by Brazelton (1). NBAS is based on the concept that a newborn who is going to have problems with neurodevelopment and behavior in later life could be identified at birth by careful observation.

NBAS assesses the newborn in seven different areas; habituation, orientation, motor component, range of state, regulation of state, autonomic activity, ability and reflexes (1). With NBAS, motor maturity of a newborn is assessed by motor component and reflexes. Though NBAS is a useful tool in research, it is rarely used in clinical practice because it is complex, time consuming and requires special training to administer. These factors make it impossible to conduct NBAS during routine neonatal examination.

In contrast, the Alberta Infant Motor Scale (AIMS) is a simple and quick assessment of motor functions of a child which can be carried out with less training. AIMS is an observational measure of motor development for infants from birth to 18 months of age, based on the neuro-maturational concept and the dynamic systems theory and measures gross motor maturation of infants from birth to the age of independent walking (2, 3). It can be easily incorporated in to the routine neonatal examination process even in low middle income countries.

Unlike NBAS which is an overall assessment of the newborn, AIMS essentially assesses only motor maturation of a child. It assesses the motor response of a child in four gravitational planes: prone, supine, sitting and standing. Thus items of AIMS focus on components those contribute to development of motor skills, such as weight bearing, antigravity movements and postural alignment (3). A score is given according to the response of the newborns when placed in these positions.

Because the Alberta scale can be incorporated into many practice settings, it is a promising method for assessing neurodevelopment among newborns. Though studies with the AIMS have been conducted on older children, there is a dearth of studies among newborns. This study was carried out to compare AIMS with NBAS when administered to newborns in assessing motor maturity at birth.

## Method

### Study setting

This study was conducted at the three obstetric units of the Colombo North Teaching Hospital in Ragama; it is one of the largest tertiary care hospitals in Sri Lanka.

### Training of research assistant (RA)

Prior to the commencement of the study, a RA who was a medical graduate was given a two-week hands-on-training by a consultant paediatrician familiar with both the AIMS and NBAS. The RA performed assessments on newborns in the presence of the trainer until the trainer was confident of the ability of the RA to conduct independent assessments.

### Administration of the tools

Babies without antenatal or natal complications were recruited for the study. Administration of the developmental tools was done on the second day of life to eliminate any influences of stress of delivery in the assessment after obtaining informed written consent from the mother. A total of 66 babies underwent both assessments. As NBAS is started with the baby in the sleeping state, it was administered first. Besides the motor component, all other components of the NBAS were also administered. AIMS was administered about one hour after administering the NBAS. Every tenth assessment was carried out in the presence of the trainer, who scored independently. The findings of the trainer and the research assistant were compared to assess the reliability of findings.

Tonus, maturity, pull to sit, defense and activity were assessed as the motor cluster within NBAS. Planter grasp, Babinski sign, ankle clonus, rooting, sucking, glabella tap, passive movements arms, passive movements legs, palmer grasp, placing, standing, walking, crawling, incurvation, tonic deviation of head, nystagmus, tonic neck reflex and Moro reflex were assessed and considered as components of motor development. Scoring was done according to the guide given in the NBAS manual (1). AIMS was administered by observing the baby in prone, supine, sitting and standing positions and observing the response. Scoring was done according to AIMS guidelines.

### Ethical issues

Approval to conduct the study was obtained from the Ethics Review Committee of the Faculty of Medicine, University of Kelaniya. Permission to conduct the study was obtained from the Director of the Colombo North Teaching Hospital, Ragama and from the Ethics Review Committee of the University of Alabama at Birmingham. Informed written consent was obtained from mothers before recruiting their babies for the study. Hand washing was done before examining babies. Examination was terminated if a baby became unduly distressed during the examination. If the baby had a deficit, the baby was referred to a Paediatrician in the same hospital for advice.

## Results

Selected family and child characteristics are summarized in Table 1. There was almost an equal number of boys and girls in the sample.

**Table 1.** Selected family and child characteristics.

| Variable | Values |
| --- | --- |
| **Maternal age** |  |
| Mean maternal age | 27.4 ± 3.9 years |
| Range of maternal age | 19.0 – 34.0 years |
| **Maternal education** | **Number (percent)** |
| Primary(grade 1-5) | 0 (0%) |
| Secondary(Grade 6-12) | 49 (75.4%) |
| Vocational and Higher education | 16 (24.6%) |
| **Paternal education** |  |
| Primary(Grade 1-5) | 0 (0%) |
| Secondary(Grade 6-12) | 50 (76.9%) |
| Vocational and Higher education | 15 (23.1%) |
| **Family income** |  |
| <20,000 Sri Lanka rupees | 32 (48.5%) |
| ≥20,000 Sri Lanka rupees | 34 (51.5%) |
| **Ethnicity of mother** |  |
| Sinhalese | 63 (95.5%) |
| Sri Lankan Tamil | 3 (4.5%) |
| **Ethnicity of father** |  |
| Sinhalese | 62 (93.9%) |
| Sri Lankan Tamil | 3 (4.5%) |
| Moor | 1 (3.0%) |
| **Birth order** |  |
| First born | 28 (42.4%) |
| Para >1 | 38 (57.6%) |
| Birth interval < 2 years (n=38) | 9 (23.7) |
| **Gender** |  |
| Male | 34 (51.5%) |
| Female | 32 (48.5%) |
| **Mode of delivery** |  |
| Vaginally | 50 (75.8%) |
| Caesarian section | 16 (24.2%) |
| **Maturity** |  |
| Mean gestational age at birth in days | 273.43±7.59 |
| Preterm | 3 (4.5%) |
| **Growth Parameters at Birth** |  |
| Mean Birth weight | 3.052 ±0.44 Kg |
| Low birth weight babies | 6 (9.1%) |
| Mean Birth length and SD | 50.53 cm ±2.23 |
| Mean Head circumference and SD | 33.82 cm ±1.48 |
| **Apgar Scores** |  |
| Mean Apgar at 1 minute and SD | 9.58±0.95 |
| Mean Apgar at 5 minute and SD | 9.96±0.27 |
| Mean Apgar at 10 minute and SD | 9.99±0.12 |

The average maternal age was 27.4 years. None of the mothers had any chronic or infectious disease during the pregnancy. Both parents of all children were educated beyond primary school and the monthly family income in the majority was more than Sri Lanka Rupees (SLR) 20,000 (1 USD ≈ SLR 150). The majority ethnic group of parents of children was Sinhalese. Among 38 children who were not the first born, the birth interval between the index child and the immediately elder sibling was less than 2 years in 9 children (23.7%).

Fifty babies (76%) were delivered vaginally. The gestational age at birth was 39.1 weeks. Only 3 babies (4.5%) were premature and only six babies (9.1%) had low birth weight. As indicated by the Apgar scores, none of the babies had birth asphyxia or needed neonatal intensive care.

Irrespective of maturity, sex, birth weight or socio-demographic characteristics, all babies had scored on the 75^th^ percentile in the AIMS. In the NBAS, there was a significant variation in the Brazelton motor score scaled to 100(figure 1). Low birth weight babies showed a narrower variation in the NABS score (figure 2). None of the scales indicated a motor deficit in any of the children.

**Figure 1.**
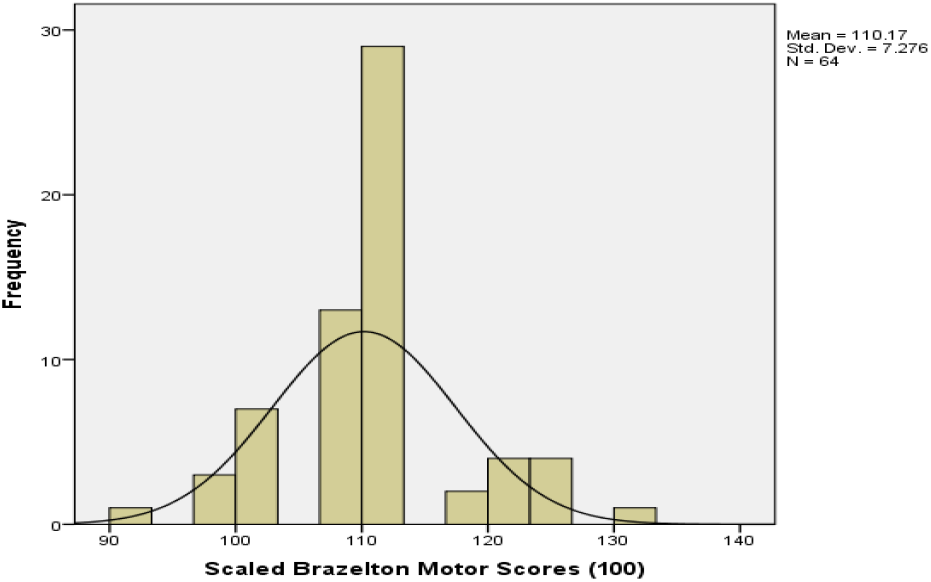
NBAS for total sample

**Figure 2.**
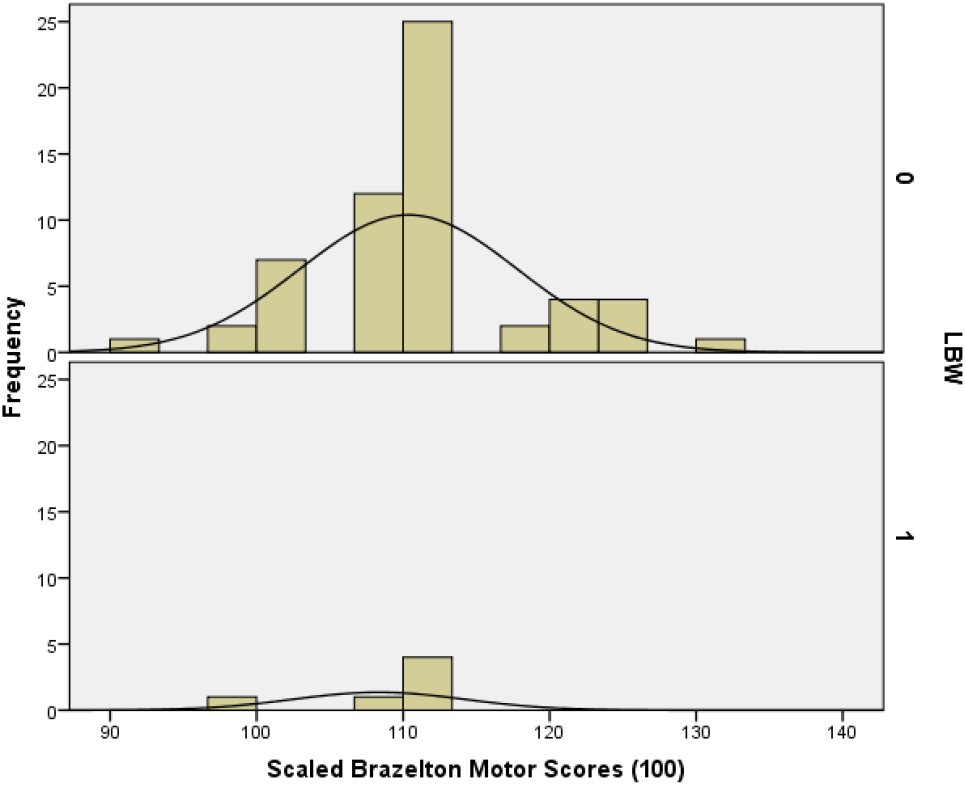
NBAS score of normal weight babies at top and LBW babies at bottom

## Discussion

Neurodevelopment is the process through which a child learns living skills to survive independently. Over the years several tools have been developed to assess and monitor neurodevelopment of children. Though clinicians are familiar with neurodevelopmental tools/scales like Bayley, Griffith and Denver, many are unaware of the Brazelton NBAS and Alberta infant motor scale. Recognition of a child with a motor problem in the neonatal period will help to improve the overall outcome if a plan of management and appropriate interventions are instituted early, resulting in reduced disability and complications. Often conditions like cerebral palsy are detected several months after birth. In some instances, it is the mother who seeks medical attention because of her concerns about motor development of the child. Hence, by the time active interventions are implemented the child is already having complications.

Though a routine neonatal examination is carried out in all newborns before discharge from hospitals, a formal motor maturity assessment is not routinely carried out. This is partly due to lack of awareness among paediatricians about availability of motor maturity assessment tools such as the NBAS and the AIMS. Even when paediatricians are aware of such assessment tools, administering the NBAS on all newborns is not practical as it is time consuming, and needs assessment items and trained personnel.

The AIMS is a simple tool to administer and does not require any special objects for assessment. A nurse or a therapist can be trained to administer the AMS which takes only a few minutes. AIMS is a good tool to screen for motor development in resource-limited settings even in busy ward settings in developing countries such as in Sri Lanka because of its practicality and psychometric properties; internal consistency of motor criteria.

The main object of this study was to compare the performance of AIMS and NBAS. Characteristically a screening tool should have a high sensitivity, where it is capable of picking up almost all positive cases. Many studies have reported that AIMS is routinely used to detect older children with motor problems early; however, we did not find any studies done on neonates (4, 5, 6).

Pai-jun et al (2004) showed that AIMS has a ceiling effect and measures infant ability best from three to nine months of age (7).Jeng et al, (2000) concluded that measurements obtained using AIMS have acceptable reliability and concurrent validity, but had limited predictive value for evaluating preterm Taiwanese infants (8).

In our study all the term neonates who were administered AIMS had no variation in the scores implying no differences in their motor maturity. However NBAS administered to the same cohort of babies demonstrated a wide variation in the motor components of the scale, although no frank defect was detected in any of the neonates. The variation in the NBAS scores indicate that differentials in motor maturity of these neonates exist which were not captured by the AIMS probably due to the small number of items available to test motor functions in the AIMS as well as scores for each item having a very narrow range. In comparison, NBAS has five motor items with a wider range of scores and 18 reflexes.

Our study sample consisted only of term babies with no antenatal or natal complications, so that difference between babies would have been too subtle to be detected by AIMS. It will be interesting to see how AIMS performs in a cohort of babies with antenatal/natal complications and normal term babies and its ability to predict future outcomes.

## Conclusion

NBAS identifies subtle differences in motor maturity of full term babies that the AIMS fails to detect.

## Acknowledgments

We thank the mothers who were willing to participate in the study.

## References

1 Brazelton T B, Nugent J K, Clinics in Development Medicine No 137 Neonatal behavioral assessment scale. 3^rd^ edition. 1995. Cambridge University Press.

2 Piper MC, Pinnell LE, Darrah J, et al. Construction and validation of the Alberta Infant Motor Scale(AMS). Can J Public Health. 1992:83(suppl 2):S46-S50.

3 Piper MC, Darrah J, Motor assessment of the developing infant. Philadelphia, Pa:WB Saunders Co; 1994.

4 Motor development of preterm infants assessed by the Alberta Infant Motor Scale: systematic review article. Fuentefria RDN, Silveira RC, Procianoy RS.J Pediatr (Rio J). 2017 Jul - Aug; 93(4):328-342. Epub 2017 May 12

5 Concurrent validity and reliability of the Alberta Infant Motor Scale in infants at dual risk for motor delays. Snyder P, Eason JM, Philibert D, Ridgway A, McCaughey T.PhysOccupTherPediatr. 2008; 28(3):267-82.

6 de Albuquerque PL, Lemos A, Guerra MQ, Eickmann SH. Accuracy of the Alberta Infant Motor Scale (AIMS) to detect developmental delay of gross motor skills in preterm infants: a systematic review. DevNeurorehabil. 2015 Feb; 18(1):15-21. Epub 2014 Oct 3.

7 Pai-jun M L, Campbell. Examination of the item structure of the Alberta Infant Motor Scale. Pediatric Physical therapy. 2004. Department of Physical therapy, university of Illinois, Chicago.

8 Jen SF, Yau KIT, Chen LC, Hsiao SF. Alberta Infant Motor Scale: reliability and validity when used on preterm infants in Taiwan. PhysTher. 2000;80:168-178

